# IMMUND: an Early Stage Diagnostic and Therapeutic Frame for Neurodegenerative Diseases and Multiple Sclerosis based on Immunological Markers

**DOI:** 10.1101/2021.05.09.442890

**Authors:** Sagnik Sen, Ashmita Dey, Dwipanjan Sanyal, Krishnananda Chattopadhyay, Ujjwal Maulik

**Affiliations:** Computer Science and Engineering, Jadavpur University, Kolkata, India; Raja S.C. Mullick Road, CSIR-Indian Institute of Chemical Biology, Kolkata, India

## Abstract

For neurodegenerative diseases, the impact of immunological markers is one of the modern research areas. It has been observed that neuroinflammation increases the cellular precipitation of some of the key proteins associated with neurodegenerative diseases. Therefore, the possibility of functional loss can be enhanced due to neuroinflammation which leads to the initiation of any related diseases. In this regard, autoantibodies, which are known for their autophagy nature, can be considered as key elements for early diagnostic as well as early therapeutics. In this article, we have proposed a comprehensive framework to unveil the diagnostic as well as the therapeutic possibility of the autoantibodies which are largely associated with Mild-Moderate Alzheimer’s Disease, Early-Stage Parkinson’s Disease, and Multiple Sclerosis. Here, we have introduced a new concept of average p-value where multiple p-values of an autoantibody in a singular disease have been considered as a multi-occurrence of that sample in cellular systems. Also, multiple proteins from a single protein family under a differentially expressed range have been prioritized. As a result, the top ten autoantibodies have been selected for further study and also considered as diagnostic markers. Interestingly, most of the selected autoantibodies are either cytokines or immunoglobulins. Subsequently, we have performed an evolutionary sequence-structure space study to identify the druggable structural facet for the selected autoantibodies. To make the therapeutic perspective more robust, we have introduced the concept of protein moonlighting. Hence, it provides more robustness in therapeutic identification. Finally, two autoantibodies i.e., Q9NYV4 and P01602 are identified as a novel marker.

## Introduction

Immune dysregulation, which refers to the breakdown in molecular control of immune system processes, is believed to be involved in neurodegenerative diseases (NDs). In case of NDs, microglia plays a vital role^1^. Microglial cells are a specialized population of macrophages in the central nervous system (CNS). It has already been reported how the NDs might be propagated through active immunophenotype^2,3^. Microglia represents a stable and inactive immunophenotype in a healthy brain, which can control the regulation of synaptic plasticity in the steady state^4^. However, microglial activates the complex immune response by increasing the expression of the pro-inflammatory cytokines and Toll Like receptors (TLRs)^5^. A strong inflammatory response has been propagated through adaptive immune cells while observing intrusions of pathogens or any chronic injury^6^. During the process, blood based immune cells like T-cell or B-cells are strongly involved^7^. The activation of adaptive immune response has also been observed in autoimmune diseases like Multiple Sclerosis (MS)^8^. On the other hand, a strong neuro-inflammatory signal may be triggered during misfolded protein deposition for NDs^9^. Although the relationship between immune responses and NDs is not simple and straight forward, in most of the cases, affective involvement of the immune responses has been observed in the later part of the disease processes^10^. In Alzheimer’s Disease (AD), the prime reason is the misfolding of the amyloid-beta (A*β*) and deposition of aggregates^11^. This soluble A*β* is responsible for the dysfunctional activations of microglial cells by interacting with cytokines, receptors, and pro-inflammatory mediators^12^. Similarly, the microglial are associated with the phagocytosis of extracellular A*β* degradations and formation of A*β* fibrils^13^. Systematic infections have been reported to be responsible for the increase in the possibility of AD by increasing the expression of TLRs^14^. Also, the inflammatory modifications were already observed in Mild Cognitive Impairment (MCI) stages^15^. However, it is difficult for the microglial cells to produce the inflammatory/ immune based fatal signals after early phase outburst of the microglial cells^16^. Therefore, the transition stage from MCI to AD is ideal for early stages diagnosis of the disease^17^). Parkinson’s Disease (PD), the second most common ND, involves the initial deposition of misfolded alpha-synuclein in dopaminergic neurons at Substantial Nigra (SN)^18^. The pathological evidences of the PD cases have shown a similar type of chronic neuroinflammation and microglial cell activation^19^. In this case, the exceeding amount of TLRs, ILs, and chemokines have been in disease cases comparing with the healthy brains^20,21^. All these results support the fact that innate immunity has a clear contribution towards the pathogenesis of NDs. Although there is no promising therapeutic intervention, some anti-inflammatory drugs have been proposed to delay AD progression^22^. Interestingly, the autoantibodies (Aabs) have been observed as potential biomarkers for NDs and autoimmune diseases like MS. These potential biomarkers were reported to be involved in the activation of the dysfunctional Microglial cells at very early stages. Therefore, Aabs are considered as influential candidates for the early diagnosis as well as early therapeutic^23^. Innate and adaptive immune responses are associated with different set of NDs. In the case of NDs, the misfolded and aggregated candidate proteins bind with the neuroglia^24^. For example, the activated microglia bind with amyloid-beta and tau protein in a multi-faceted manner. All functions have been characterized by cytokines, TLRs, TNFs, etc. Immuno-precipitations are highly associated with the loss of cognitive responses. In contrast, MS is an autoimmune disease, and hence, the involvement of the Aabs is highly significant in MS. It has been observed that the early stages of different NDs (like AD and/or PD) and MS might share same set of Aabs (Give references). However, the Aabs should be functionally segregated in different diseases. This hypothesis can provide the possibility of using Aabs for the diagnosis as well as therapeutic applications against these categories of neuronal dysfunction. In this article, we aim to investigate potential autoantibodies for early detection of the AD, PD, and MS. We have started with understanding differential expression of the autoantibodies for Early Stage Parkinson’s Disease (ESPD), Mild Moderate Alzheimer’s Disease, and MS where MCI has considered as a controlled stage. Only the common samples were considered for further study. The list of autoantibodies has been filtered based on few parameters. The final samples can be considered as diagnosing biomarkers for these early stages. Subsequently, these samples have been studied based on their structural properties such as disorder predictions, sequential stability index, evolutionary sequence space analysis. The sequence information was then mapped on individual structure networks using an Antistrophic Network Model. This process helps to unveil the structural cores that can be targeted by potential drug candidates. Finally, the specific sets of pathways have been observed to understand the possibility of mutually exclusive functional duality for each candidate.

## Results

In this study, we used a combination of different approaches including graph theory, evolutionary analyses and structural studies (please refer to the flowchart in Figure 5) to investigate the common biomarkers of Alzheimer’s (MMAD), Parkinson’s disease (ESPD) and multiple sclerosis (MS). Different components of this flowchart have been discussed in detail in the Methods section. For this study, we have used protein microarray data from GSE74763 dataset^25^ containing 50 mild cognitive impairment (MCI) samples and 50 disease samples (curated from NCBI). These data samples were further investigated for differentially expressed auto antibodies biomarkers which could be utilized as diagnostic indicators for these diseases.

**Figure 1.**
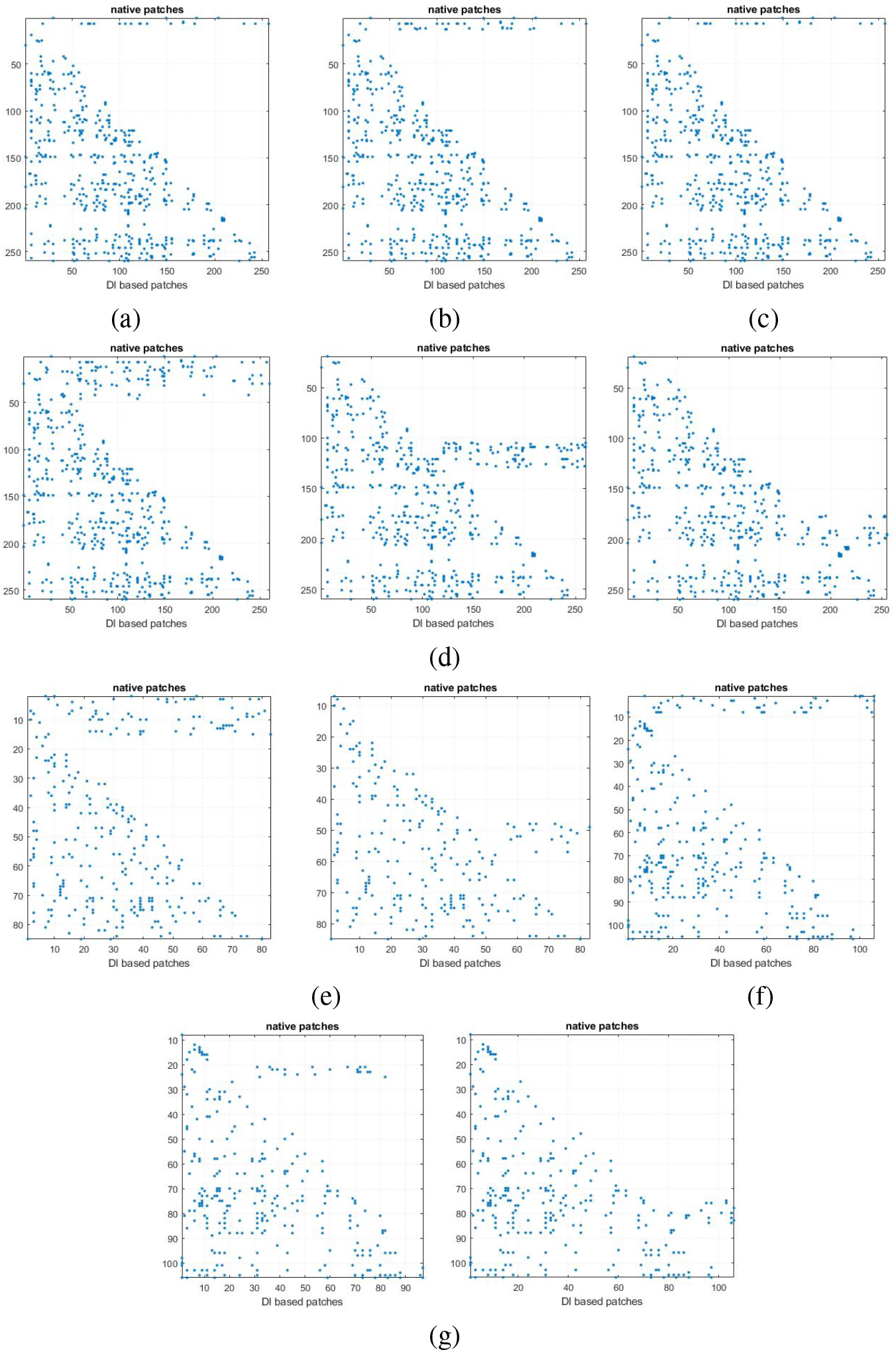
Direct Coupling Analysis of the amino-acid pairs directly involved in evolution of proteins (a) Phosphorylase b kinase gamma catalytic chain, testis/liver isoform, (b) Serine/threonine-protein kinase 36, (c) Serine/threonine-protein kinase PAK 1, (d) Cell division cycle 2-related protein kinase 7, (e) immunoglobulin lambda locus (f) immunoglobulin kappa variable 1-5 and (g) immunoglobulin lambda variable 2-14.

**Figure 2.**
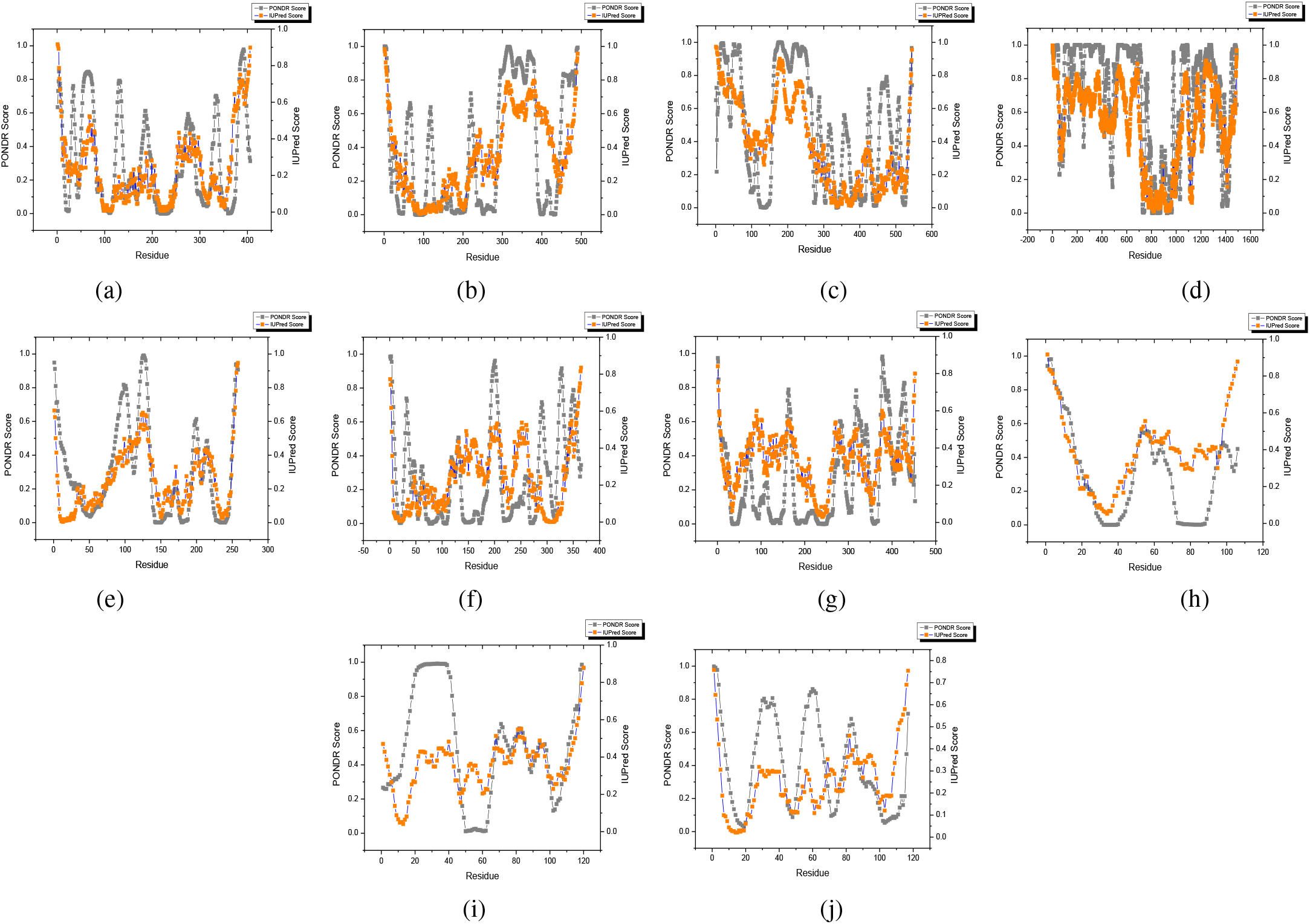
The disorderness of the proteins (a) Phosphorylase b kinase gamma catalytic chain, testis/liver isoform, (b)Serine/threonine-protein kinase 36, (c) Serine/threonine-protein kinase PAK 1 and (d) Cell division cycle 2-related protein kinase 7 (e) Ig lambda chain C regions, (f) Fc fragment of IgG, low affinity IIIa, receptor, (g) Immunoglobulin heavy constant mu (h) Immunoglobulin lambda locus (i) Immunoglobulin lambda variable 2-14 and (j) Immunoglobulin kappa variable 1-5 are predicted and represented the score of IUPred and PONDR prediction model through orange and grey lines respectively.

**Figure 3.**
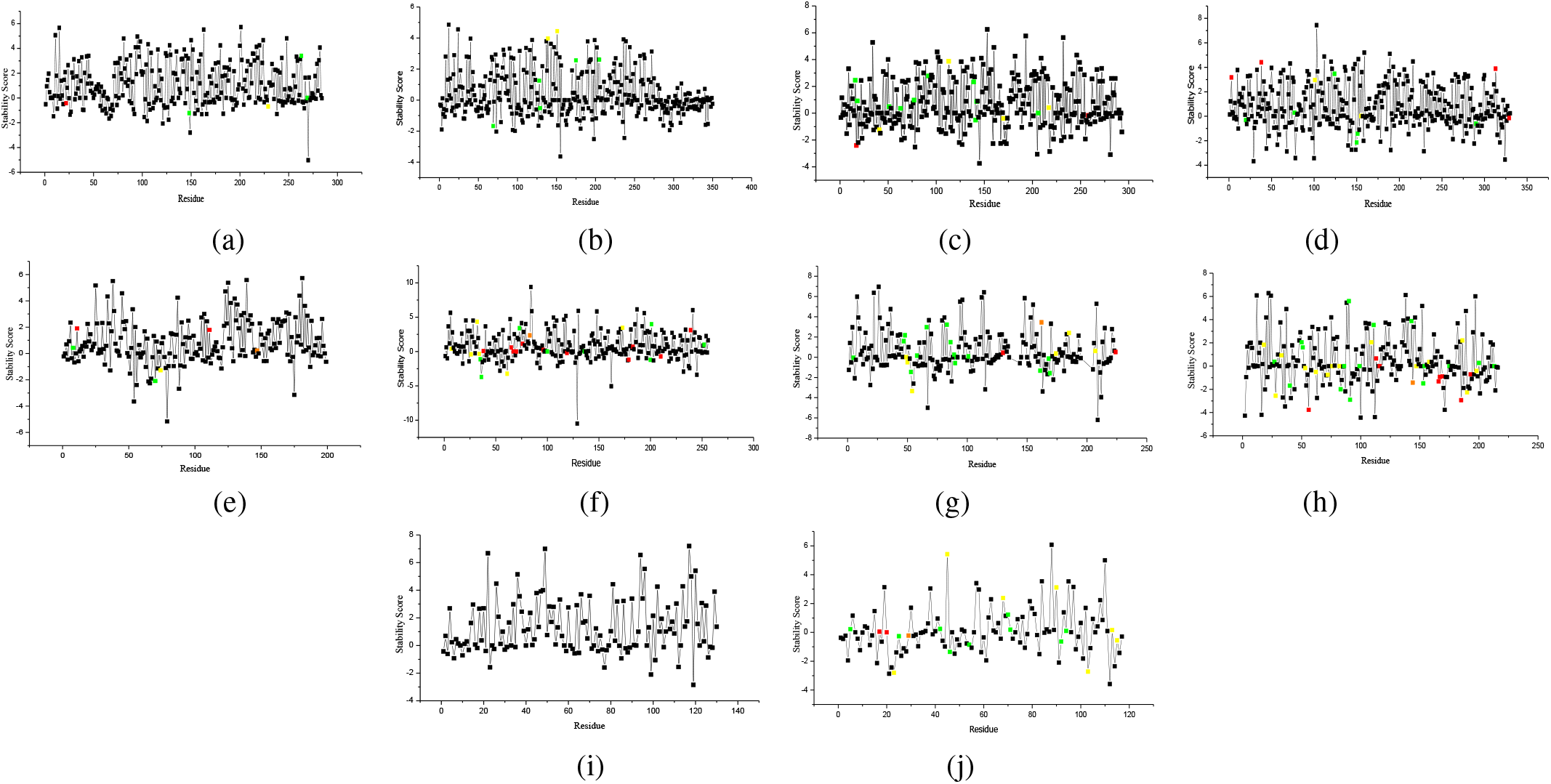
Alascan mutagenesis of the proteins (a) Phosphorylase b kinase gamma catalytic chain, testis/liver isoform, (b)Serine/threonine-protein kinase 36, (c) Serine/threonine-protein kinase PAK 1 and (d) Cell division cycle 2-related protein kinase 7 (e) Ig lambda chain C regions, (f) Fc fragment of IgG, low affinity IIIa, receptor, (g) Immunoglobulin heavy constant mu (h) Immunoglobulin lambda locus (i) Immunoglobulin lambda variable 2-14 and (j) Immunoglobulin kappa variable 1-5. The functional non-bonding energies such as van der Waals, Chi-angle, Torsion angle, cis bond of each amino-acid is represented by colour green, red, yellow and orange respectively.

**Figure 4.**
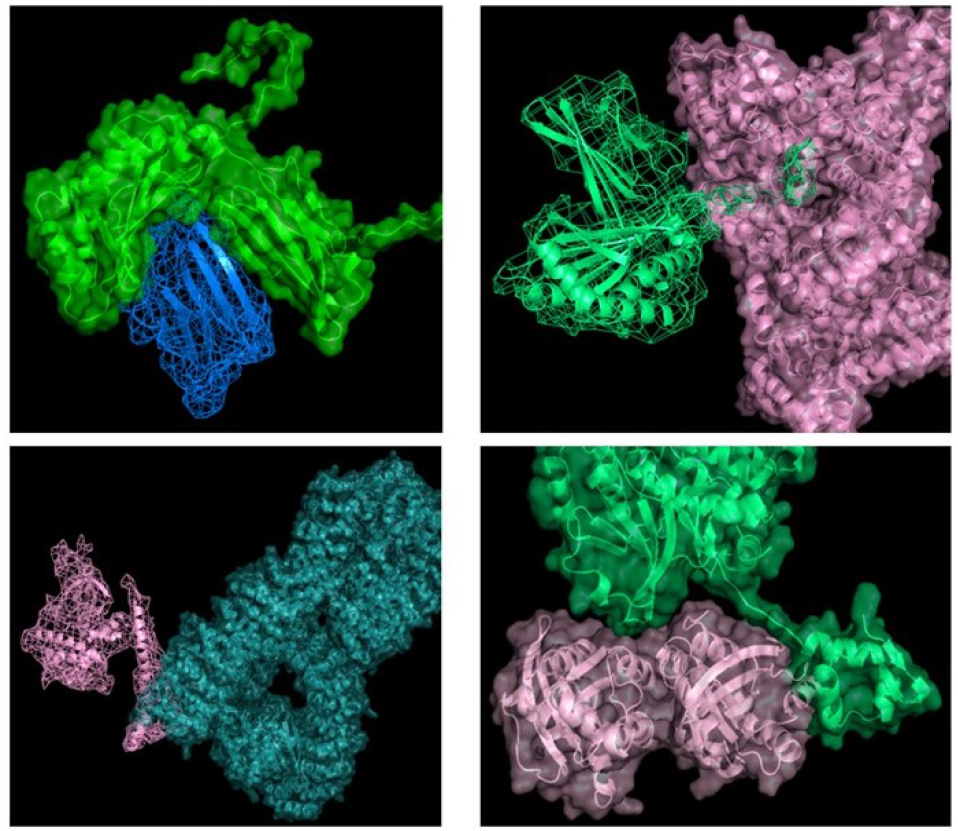
The Docking validation study and corresponding docking postures

**Figure 5.**
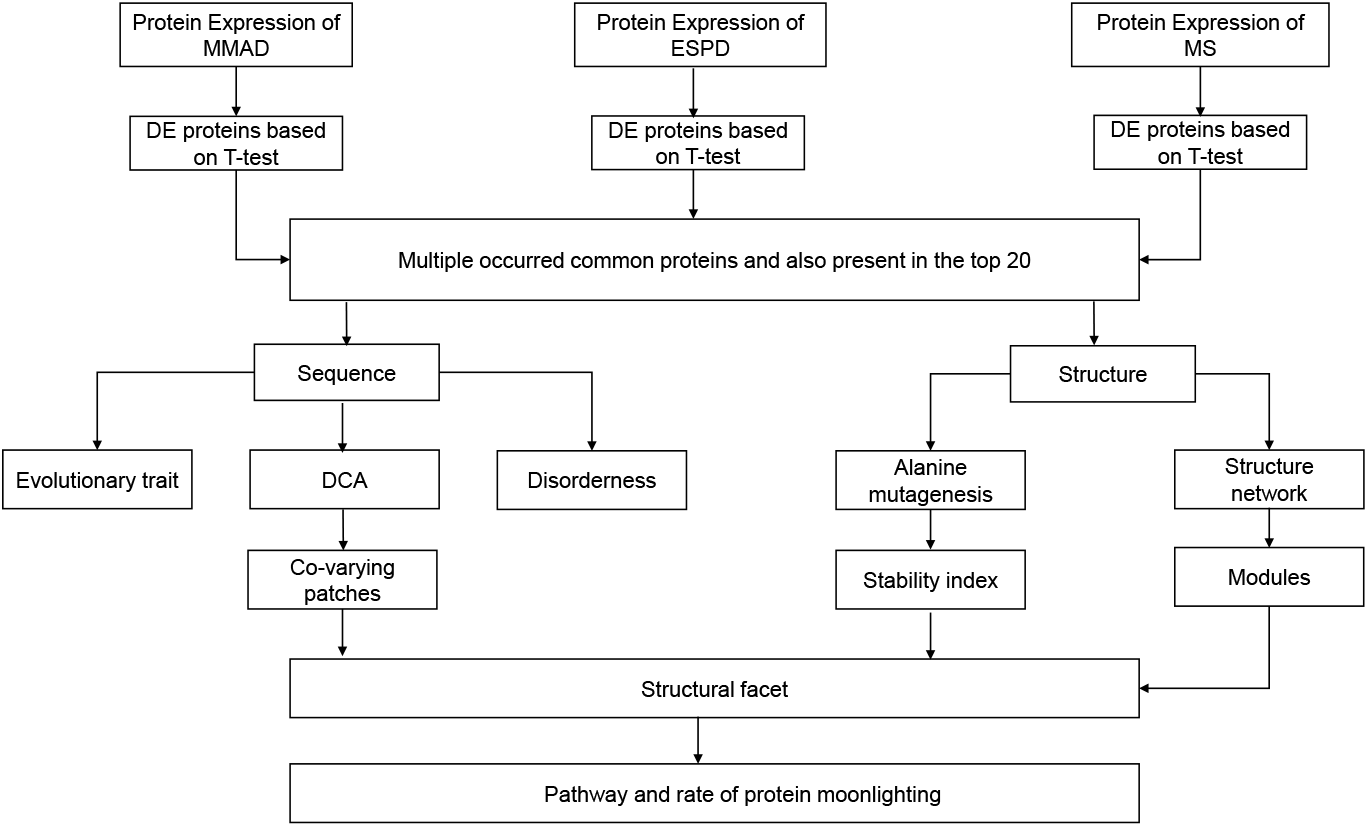
Flowchart representing the working methodology

One of the primary aims of this study is to understand the structural facet of autoantibodies common for three selected diseases. First, we performed “two sample T-tests” to determine the mean of the expression values, which can be different from the hypothesized or known population mean. The probability value of observing the t-test results under the null hypothesis helped to understand the statistical significance. This value is also known as p-values, the details of which for each protein have been given at supplementary Table T2. We considered in this study proteins with p-value less than 0.05 as the differentially expressed antibodies (DEA). After identifying the DEAs, 195 common autoantibodies of the three selected neurodegenerative diseases were selected (as shown in Figure 6). After parameterized sorting, top 10 proteins were chosen for subsequent investigation, which included the (a) evolutionary analyses, (b) structure space analyses, and (c) multiple function analyses.

**Figure 6.**
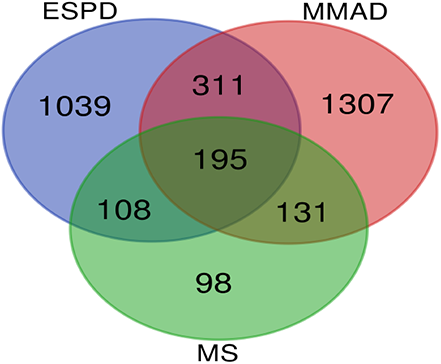
Differentially functioned autoantibodies collectively working as biomarkers for Alzheimer’s, Parkinson’s and Multiple sclerosis

### Sequence Space Analyses

Top ten selected proteins were found distributed in three families, Viz., Immunoglobulin C1-set domain (Pfam id-PF07654), Immunoglobulin V-set domain(Pfam id-PF07686), and Protein Kinase (Pfam id-PF00069). Among them, a total of three members (UniProt Id. P55899, P01871, and P0DOY3) with multiple p-value occurrences belonged to two families i.e., PF07654 and PF07686 (Supplementary Table T3). From the initial Shannon Entropy (SE) scoring, we found that these two families have a higher trait of disorderness (Table 1). In addition, these families showed multiple binding partners or multiple binding propensities with the same partner (defined by the term multi-occurrences) (Supplementary Table T3). However, in the case of the third family (Pfam ID: PF00069) the disorder trait was found to be lower (as the threshold is 2.9) (Table 1). Interestingly, the proteins with higher values of disorder trait were found belonging to the immunoglobulin family proteins (PF07654 and PF07686). We then identified that six out of the ten proteins were immunoglobulins, while the other four Aabs were kinases (belong to PF00069). Then, using the sequence information of the above mentioned three protein families we performed multiple sequence alignment (MSA) followed by direct coupling analysis (DCA) study (Figure 1). Through the course of evolution the impact of substitution of a residue at a particular site was/would be nullified by the alteration at another position in close proximity in the 3D protein structure. These two positions are called to be the evolutionary coupled/co-varying pairs. DCA study helped us identify the coupled pairs, which played a key role to understand the important region in the protein structures. Hence coupling propensities at IDRs have been computed. The disorderness of individual protein was predicted from two prediction models, namely PONDR-VLXT and IUPred-s. The regions found to be common using both algorithms were considered to be the effective disordered regions (Figure 2). The outcome was complying with the SE score obtained for the three families. The predictive disorders were mostly justified by the higher SE scores for two families (PF07686 and PF07654) and by lower SE score of P0DOY3 family proteins. Similarly, IDRs within the evolutionary coupled patches helped to explore the distribution of disorderness. More elaborately, 7 out of 10 proteins had disordered regions, found to exist within evolutionary co-varying patches Figure 1. Other three were not shown in Figure 1 due to extremely low coupling propensities. Among the seven proteins, PF00069 family has four members, PF07654 family has one member, and finally, PF07686 family has two members in the list. Out of these four multi-occurred samples, two proteins (UniProt id P0DOY3 and P01602) are shown in Figure 1e and Figure 1f, respectively. It can be suggested that the disordered regions with high co-varying propensity would have significant impact on the overall structure of the proteins. We infer that such effective disorderness can partially enforce to create globule-like structure at monomeric stage that justifies the presence of the maximum number immunoglobulin families in the disorder list. The protein members from PF00069 were found to be composed of many co-varying patches that were distributed mainly within a particular stretch or region. These have been termed in this article as locally distributed co-varying patches (Figure 1a-d). Among them, protein (uniprot id: P0DOY3) was found to have two disordered regions, one from 2 to 15 (position from 2 to 7 were considered to be effective disordered region) and another from 48 to 58 (48-54 and 57 were in the effective disordered regions). For the first disordered range, 24 unique residues were predicted to be coupling pairs with different direct information (DI) score (Supplementary Table T4). In the second disordered range, 12 different amino acid positions were observed to be co-varying. Similarly, protein P01602 was found to be composed of mainly one effective disordered region (sequence 1-8). Interestingly, the amino acid sites within these ranges were observed to couple with 35 different positions throughout the sequence. Similarly P15735, Q13153, and P01704 proteins had 21, 21 and 17 coupled pairs that were distributed mainly in the ordered regions. Here, it was observed that the positions in the proteins that belong to the families with lower disorder traits (having low SE score) mainly had coupled pairs that showed the propensity to be ordered. The aforementioned studies had provided clear insight regarding the evolutionary conservation, covariation as well as individual sequential orchestrations. However, the observation can be more stringent through structure-based study.

**Table 1.** The details information of the selected protein families

| Protein Name | Uniprot Id | PFAM Id | $P_{avg}$ | Family Trait |
| --- | --- | --- | --- | --- |
| Phosphorylase b kinase gamma catalytic chain, testis/liver isoform | P15735 | PF00069 | 5.10E-14 | 3.47 |
| Serine/threonine-protein kinase 36 | Q13188 |  | 3.07E-12 |  |
| Serine/threonine-protein kinase PAK 1 | Q13153 |  | 3.04E-11 |  |
| Cell division cycle 2-related protein kinase 7 | Q9NYV4 |  | 5.63E-12 |  |
| Ig lambda chain C regions | P04440 | PF07654 | 1.60E-12 | 4.02 |
| Fc fragment of IgG, low affinity IIIa | P55899 |  | 1.37E-06 |  |
| immunoglobulin heavy constant mu | P01871 |  | 0.004 |  |
| immunoglobulin lambda locus | P0DOY3 |  | 1.25E-05 |  |
| immunoglobulin lambda variable 2-14 | P01704 | PF07686 | 2.88E-10 | 3.73 |
| immunoglobulin kappa variable 1-5 | P01602 |  | 0.0004 |  |

### Structure Space Analysis

The structure space study has two levels of analysis: I. Structure Network Analysis, and II. Alanine Mutagenesis. Here we had compared and merged the sequence space observations with detailed information obtained from structure network study. The PDB structures of the selected protein chains that were associated with the disease systems were chosen for this study (@refer RCSB). The selected PDB structures were then subjected to structure network study. Structure network analyses provided an understanding of the internal arrangement and inter-dependency of the amino acid residues in protein structures. In this analysis, all residue networks of the proteins were generated that were further split into interlinked community cluster networks by means of Girvan-Newman algorithm. In these cluster networks, the highly interacting amino acid residues were assembled in the residual communities (Supplementary Figure F1). Using betweenness centrality calculations, the influence of a particular residue on the overall structural dynamics was then explored by considering its fluctuations profile. The structures with maximum disorderness were found to lack a diverse set of communities based on the residue-residue interaction. In terms of color modules, residues were flocked together in a minimum number of modules for 5 out of 10 proteins (UniProt id.-Q13188, P04440, P55899, P01871, and P0DOY3). Among the above mentioned 5 proteins, the last three were multi-occurred, while the rest five proteins (UniProt ids-P15735, Q13153, Q9NYV4, P01704, and P01602) have 6, 10, 10, 10 and 12 modules respectively. Therefore, the maximum number of proteins were not able to assemble in ordered structures. Subsequently, the concept of alanine mutagenesis was introduced to interpret the contribution of a single residue towards the stability of a protein by means of the stability index value. As alanine is a non-bulky amino acid, the mutagenesis at every residue can provide the rate of stability^26^. This approach was used for all the selected Aabs (Figure 3).

The quality assessments of the selected proteins were performed and respective non-bonding clashes were calculated. By applying quality assessment, the effect of Van der Waals force (Vdw), torsional angle etc. were calculated on each residue (marked in different colors) as well as the changes in energy due to non-bonding interactions were determined (Figure 3). The H-bond clashes were calculated using following two ways, i.e., Vdw clashes and changes in torsional clashes in the residues. Following the alanine scanning results, comparative rates of fluctuations were given in color dots. In the case of four multi-occurring proteins, the Vdw clashes and the changes in torsional clashes were observed at all the disordered regions. In IGHM (Uniprot id. P01871), Vdw clash was observed at residue 200 (consisting TYR) whereas at residue position 201, photo-isomeric changes of dihedral angles were observed in terms of *ω* angle. Similarly, P0DOY3 was found to have three Vdw clashes at residue position 7, 11 and 14 (consisting of residues SER, ALA, and GLN respectively) in the first disordered region whereas second disorder region had two Vdw clashes and one torsional clash at residue position 49, 50 and 52 respectively (residue PHE, ALA, and THR). At residues 39, 26 and 61 (consisting LEU, ASN, and ARG respectively) of P0DOY3, one Vdw clash and two torsional clashes were observed. On the other hand, only Q9NYV4 has one Vdw clash at residue position 124 (consisting GLN) and coupled partner residue position 77 (consisting ASP). However, Vdw clashes are observed at coupled partners of Q81WQ3, and Q13188 at residue position 204, 69 and 129 (consisting MET, TYR and, ARG respectively) respectively. Subsequently, P01602 had one Vdw clash at residue position 5 (a PRO residue). In case of all these three samples extreme changes in *χ* angles at a nearer position of the Vdw clashes were observed, except P55899, in which changes in *ω* angle had been observed. Using the steric clashes as computed above, we defined the clashes zones at the disordered regions, which would be important patches/regions for binding interactions. In addition, based on the sequence space results, the coupled pairs were also considered to be binding facets for the proteins. Using the above criteria the non-covalent interactions at the coupled partners of the disordered regions were explored.

### Protein Moonlighting and Functional Information

It is now established that a particular protein may adopt multiple structural facets resulting in functional multiplicity (protein moonlighting). We have analyzed the moonlighting property of the selected proteins to determine their association with multiple unrelated functions. We have identified the proteins having multiple distinct functions from the sample dataset. We found that some of these proteins had no experimental evidence of multi-functionality. For such proteins, their homologs were identified and the functions of the homologs were considered for the moonlighting functions. Among them only some functions were observed to have implications on the selected diseases (Table 2).

**Table 2.**
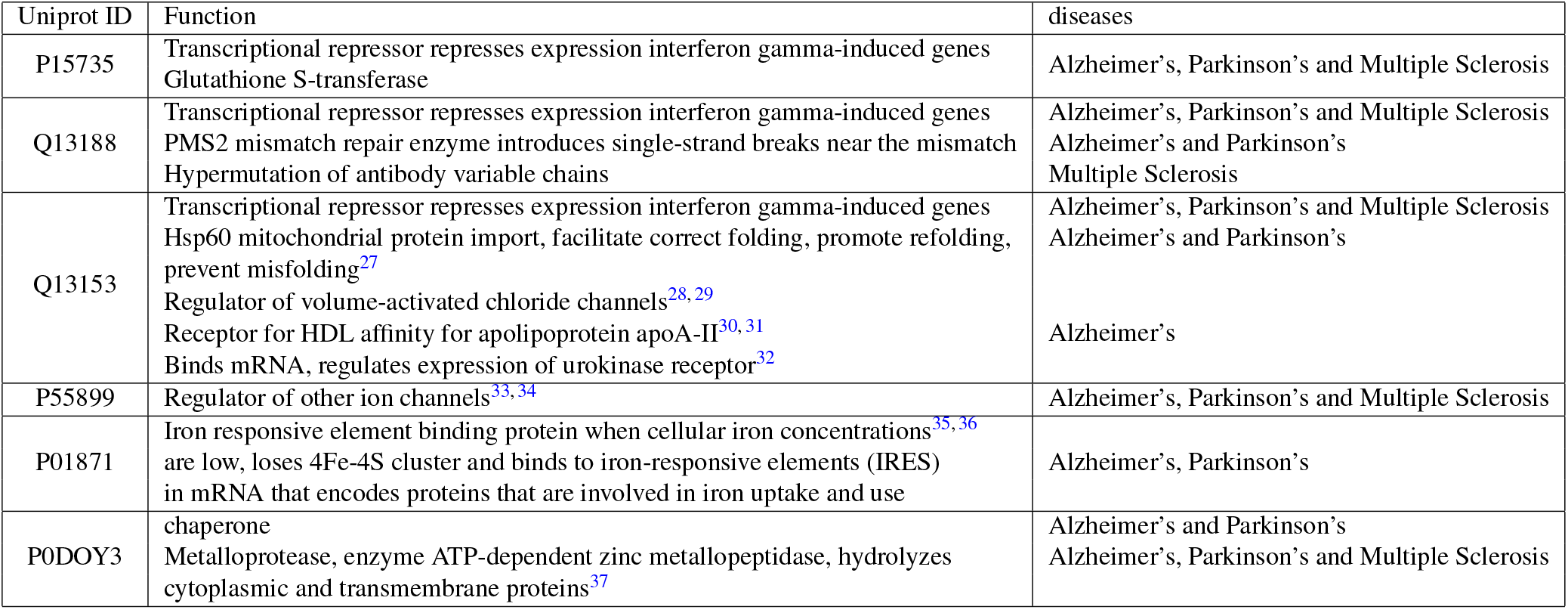
The moonlighting properties of the selected proteins

### Docking Validation

In order to validate the identified binding facets, docking study was performed with the interaction partners selected from PPI network. From protein-protein docking study, we found that 1LDS was housed in the binding groove of protein P55899. This interaction mainly involved the residue stretch from 48 to 56 of 1LDS. Since serum albumin (ALB) is also known to be a binding partner of P55899, we also used docking study to find out their interaction domains. The binding of ALB was found to occur through the terminal loop region from residue 7 to 41 of P55899. We observed that P55899 used two binding regions, which are opposite to one another, to interact with these two binding partners (1LDS and ALB). The most favorable bound conformation of Protein 2IAE and Q13188 indicated that they were in close proximity by involving 106 atoms. The terminal helix and loop regions of Q13188 were involved in this interaction, shown in Figure 4.

## Methods

In this article, an evolutionary study is performed on the common biomarkers of Alzheimer’s, Parkinson’s disease and multiple sclerosis by comparing the effects due to structural disorder. We have provided a flowchart that represents the site map of the proposed methodology in details, as shown in Figure 5.

GSE74763 dataset^25^ is used in this study which is curated from NCBI. In order to recognize the differentially expressed autoantibodies, the dataset contains 50 disease driven mild cognitive impairment samples and 50 controlled samples. These are examined onto human protein microarrays. Moreover, these samples are further studied for differenially expressed autoantibodies biomarkers which can be utilized as diagnostic indicators.

### Identification of differentially functioned autoantibodies

Two-sample t-test is applied on the samples to determine whether there is a significant difference between the means of two groups, based on the standard deviation of the groups. The significant difference between the sample data is estimated depending on the t-values. Let for each sample, group 1 consists of *N_D_* diseased sample with mean 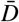 and standard deviation *S_D_* and group 2 consists of *N_C_* control sample with mean 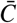 and standard deviation *S_C_*. Then, the t-value thus obtained is defined as follows:

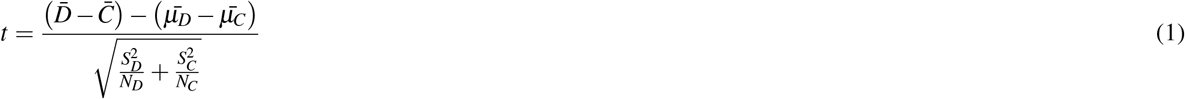

And the degree of freedom can be expressed as:

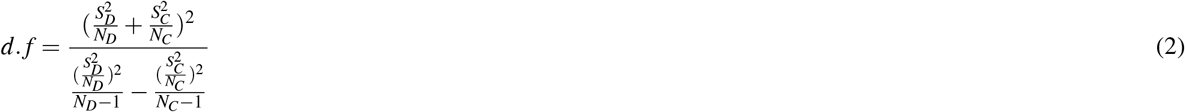

The t-test is applied on the normalized data of the three diseased dataset to identify the DEA in each case. A threshold 0.05 is considered for filtration of null hypothesis. If the p-value of a sample is less than 0.05, it signifies the protein is differentially functioned.

### Common autoantibodies determination

From the differentially expressed autoantibodies biomarkers of each disease, the common biomarkers are identified and shown through a venn diagram in Figure 6. These 195 common samples (reported in Supplementary Table T1) are further ranked based on the average p-value (*P*_(avg)_) defined in Equation 3. Here, p-values of ESPD, MMAD and MS are represented as *P* — *value_ESPD_*, *P* — *value_MMAD_* and *P* — *value_MS_* respectively and *m* is the number of diseases. However, in this research work *m* = 3.

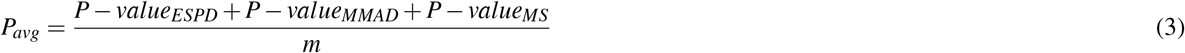

During this study, some interesting cases are observed. Several proteins occurred multiple times posses different p-values in three diseases. For those proteins *P*_(*avg*)_ is calculated in terms of special cases (*SP*_(*avg*)_), whereas the number of occurrence in a particular diseases are denoted as *n*, shown in Equation 4. These proteins are selected for further analysis as they have an important impact on three neuro-degenerative diseases.

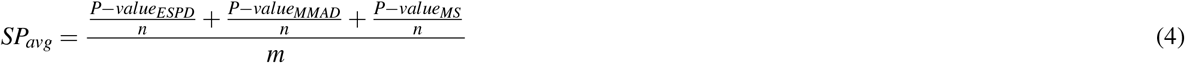

Depending on *P*_(*avg*)_ score, four proteins are selected, those are present in top 20 and also belong to a same family PF00069. In parallel, other six proteins are selected based on the (*SP*_(*avg*)_) score and they belong to PF07654 and PF07686. Strikingly, these two families are also present in top 20 rank list.

### Shannon Entropy

In order to understand the family trait of the selected three families, Shannon entropy is performed for each consensus sequence of the protein family as well as the proteins those are finally selected and belong to these families. It is known that, entropy posses an idea of disorder, directly proportional to the rate of disorder i.e. if the disorder increases it signifies higher entropy. Shannon entropy is defined as follows:

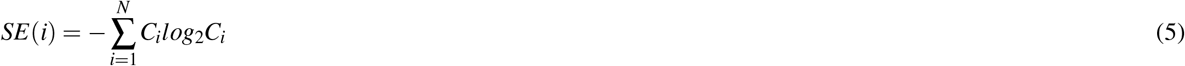

where *C_i_* was the probability of given amino acids and N was the number of letters in a sequence. The summation run over the 20 residues that normally were present in a protein sequence. The probability *C_i_* represent the composition of the consensus sequence. So the entropy range lied between 0 and the *log*_2_(20) = 4.32. Moreover, our result is mapped to Ponder and IUPred for better evaluation. From the two aforementioned tool the common coincide disorder regions are selected.

### Direct Coupling Analysis

Secondary correlation between non-interacting residues may arise from correlations between substitution patterns of the interacting ones. In order to investigate native contacts in a more specific way, direct couplings were needed to be understood explicitly. A major shortcoming of the covariance study; i.e., the MI theory, was that it cannot disentangle direct correlations from indirect ones. Direct information (DI) is one such case on MI (Mutual Information), which aims at disentangling direct interactions from indirect ones. Based on DI scoring (Supplementary Table T4), direct coupling analysis (DCA) was performed. Therefore, from DCA, a lucid view of DI (Equation6); i.e., how directly the selected coincide sites are coupled with each-other was obtained.

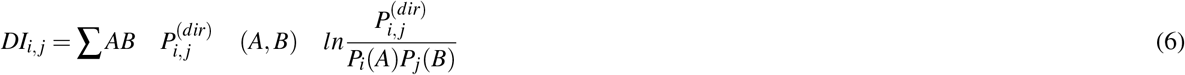

Here, Here *P_i_j*^(^(*dir*)) represents reweighted frequency counts to introduce two residues for DI. where *P*(*A*,*B*) is considered as joint probability, and *P*(*A*) and *P*(*B*) are individual probability. *P_i_*(*A*) and *P_j_*(*B*) are for amino acid type A at *ith* position and similarly B at *jth* position.

### Model Building

Here sequences of P55889 and Q13188 were retrieved from UniProt and their models were generated using I-TASSER (Zhang). Coordinate information of the partner proteins were obtained from the Protein Data Bank (PDB) (PDB ID:). The generated models and the selected PDBs were further subjected to structure network analysis.

### Structure-network analysis

In order to explore the internal organization and inter-dependency of the amino acid residues in terms of pairwise interactions in the model structures, structure network analysis was deployed. The interaction established between the elements of the networks, i.e. the residues depending on the interaction energy and/or spatial distance, were represented through edges and nodes. Reckoning on the interaction strength the sides were drawn between the two amino acid nodes. The equation was represented below.

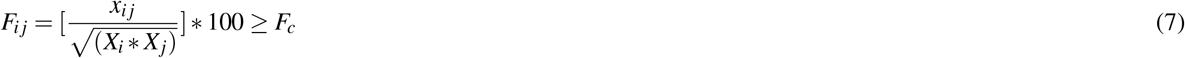

*F_c_* was the threshold of interaction strength, the default value is 4%. Here, *x_ij_* was the number of side chain atom pairs of residues *i* and *j*. *X_i_*, and *X_j_* were the normalization factor for residues types *i* and *j*^38,39^. Stability Score Calculation Additionally, stability score of the amino acid present in a protein was calculated by alascan mutagenesis technique. Similarly, steric clashes of the amino acid was also performed. This was one of the artifacts prevalent in low-resolution structures and homology models. Steric clashes arised due to the unnatural overlap of any two nonbonding atoms in a protein structure. Usually, removal of severe steric clashes in some structures is challenging since many existing refinement programs did not accept structures with severe steric clashes. Due to non-bulky, chemically inert, methyl functional group that nevertheless mimics the secondary structure preferences alanine is used to check the stability.

### Molecular docking Validation

To have mechanistic understanding into protein-protein association that has been manifested from our previous analysis, we deployed the computational strategy of protein-protein docking^40^. Models built from I-TASSER along with the selected co-ordinate structures were subjected to the docking study and tools from ClusPro were used^41^. In this rigid body docking, the angular step size for rotational sampling of ligand orientations was set to about 5° in terms of Eular angles. Energy minimization followed by root-mean-square deviation (RMSD) based clustering was performed for accurate and near-native conformational sampling and refinement of the complex structure. In each case, docked conformation having the lowest energy value was considered. Along with the energy value, conformations with the highest number of members involved were also prioritized in order to study a precise complex encounter and also to confirm that the interaction between two candidate structures took place in the neighborhood of the native state.

## Notes

### Competing Interest Statement

The authors have declared no competing interest.

