## Supplementary Figure S1 for "IMMUND: an Early Stage Diagnostic and Therapeutic Frame for Neurodegenerative Diseases and Multiple Sclerosis based on Immunological Markers"

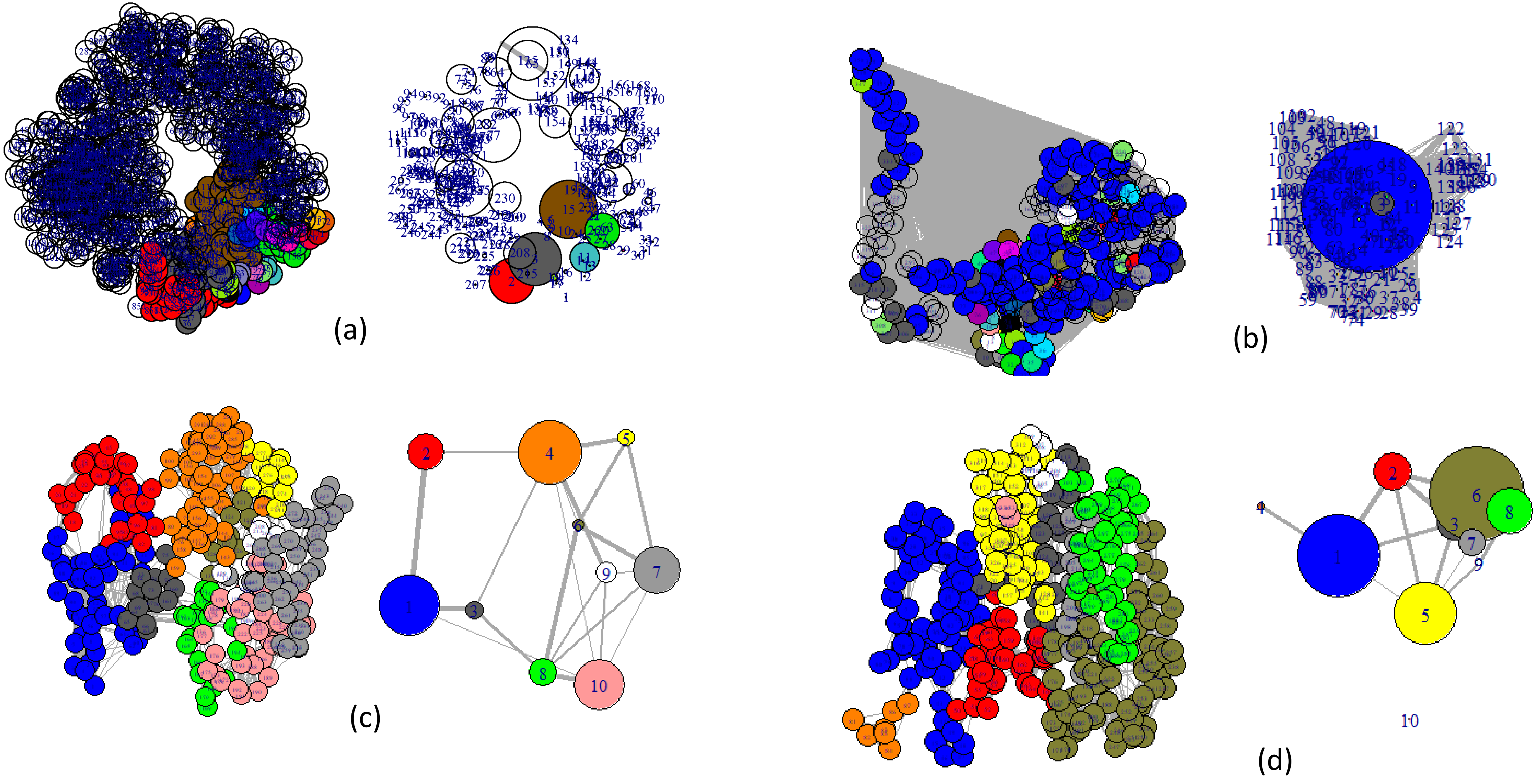


Structure network for the proteins (a) Phosphorylase b kinase gamma catalytic chain, testis/liver isoform, (b)Serine/threonine-protein kinase 36, (c) Serine/threonine-protein kinase PAK 1 and (d) Cell division cycle 2-related protein kinase 7 belong to the same family. The PFam Id is PF00069

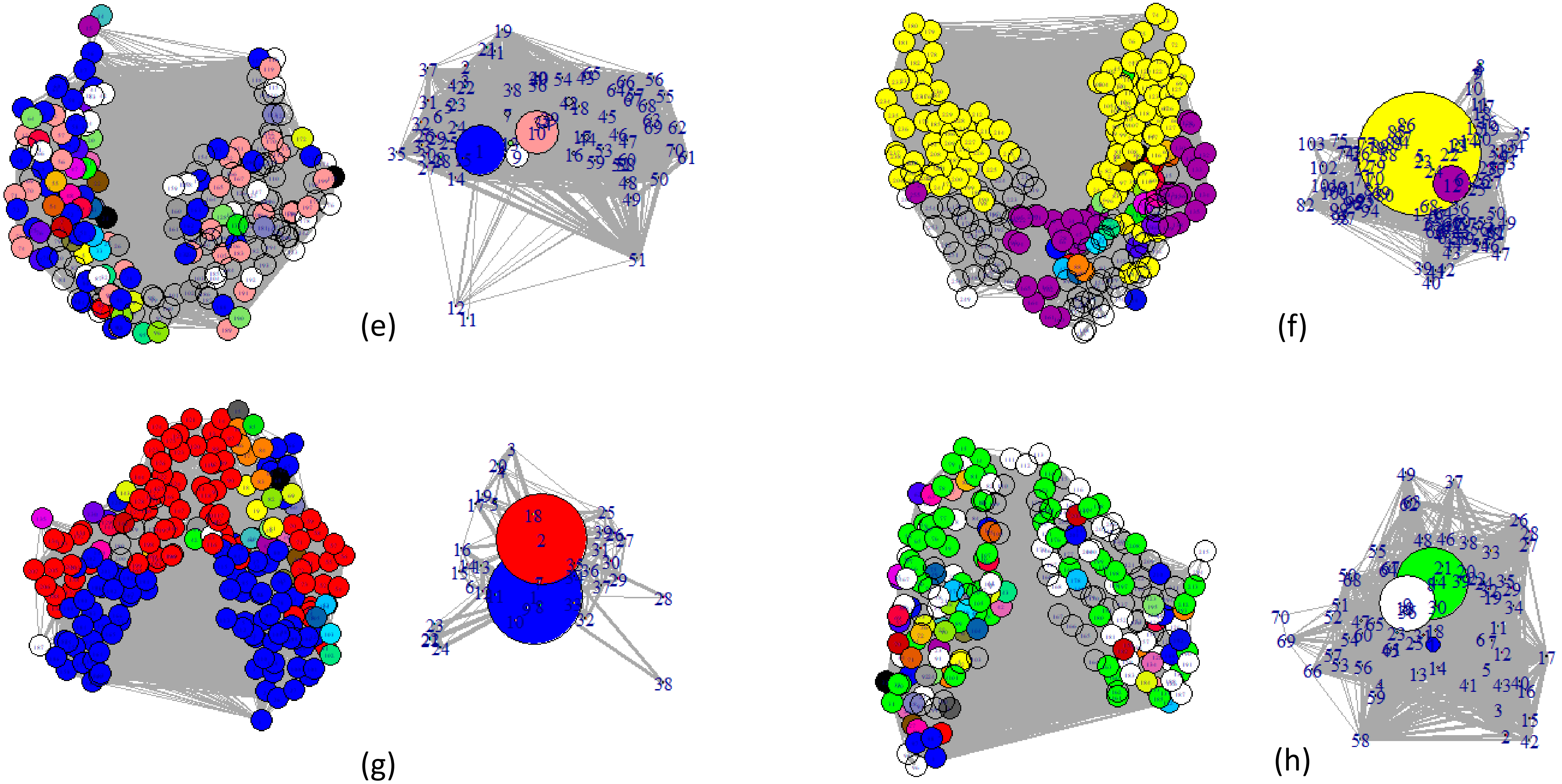


Structure network for the proteins (e) Ig lambda chain C regions, (f) Fc fragment of IgG, low affinity IIIa, receptor, (g) Immunoglobulin heavy constant mu and (h) Immunoglobulin lambda locus belong to the same family. The PFam Id is PF07654

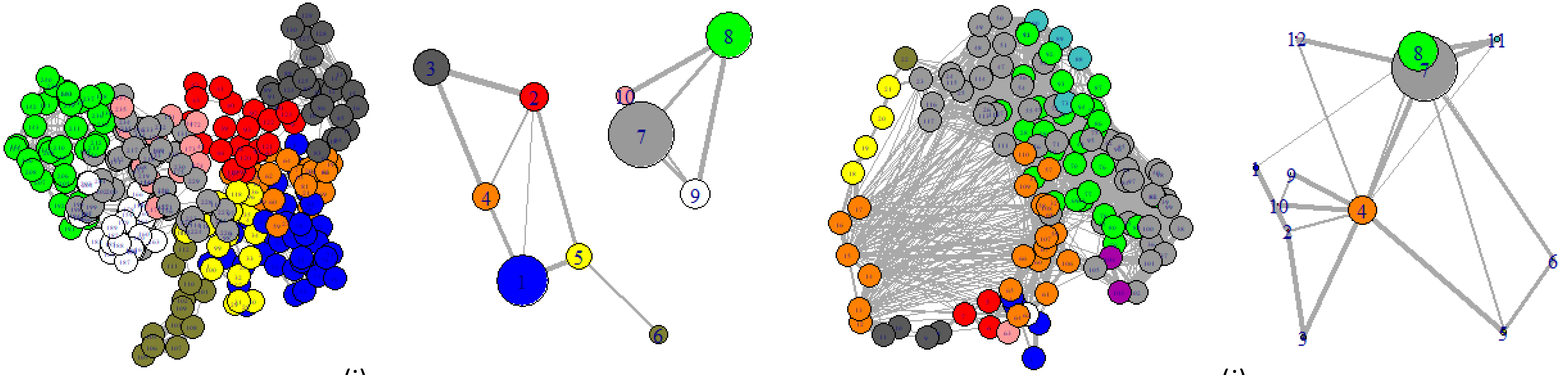


(i) (j)

Structure network for the proteins (i) Immunoglobulin lambda variable 2-14 and (j) Immunoglobulin kappa variable 1-5 belong to the same family. The PFam Id is PF07686.
